# Two-pore channel 2 is a key regulator of adipocyte differentiation via the cAMP signaling pathway with calpain as downstream effector

**DOI:** 10.1101/2021.03.20.436264

**Authors:** Yuxuan Zhang, Lai-Hing Chan, Ruth Tunn, Margarida Ruas, David Gay, Marijana Todorcevic, Costas Christodoulides, John Parrington

## Abstract

We investigated whether the endolysosomal two-pore channel TPC2 is a mediator of adipocyte differentiation. We show that *Tpcn2* mRNA is expressed transiently during induction of C3H10T1/2 mesenchymal stem cells to differentiate into adipocytes, and that this expression is triggered by cAMP. This is the first demonstration of a cell signaling pathway that can regulate TPC gene expression. We also identified an important functional role for TPC2 in adipocyte differentiation. First, ectopic TPC2 expression in C3H10T1/2 cells partially rescued the block to adipocyte differentiation caused by cAMP absence. Second, inhibition of endogenous TPC2 expression in primary preadipocytes substantially reduced their ability to differentiate into adipocytes. Finally, genetic variation at the *Tpcn2* locus is associated with increased upper-body fat distribution in women concomitant with reduced *Tpcn2* expression in abdominal adipose tissue. Our findings implicate TPC2 as an important mediator of adipogenesis and may aid identification of new drug targets for treatment of obesity.

## Introduction

The rise of obesity in the developed and developing world and the increased risk of type 2 diabetes, coronary heart disease, stroke, and cancer that accompanies this condition, is leading to increased interest in adipogenesis, the process by which adipose progenitors differentiate into mature adipocytes (Sarjeant & Stephens, 2012). Adipose tissue is a dynamic organ composed of adipocytes which are the predominant cell type, and a variety of other cell types such as preadipocytes, endothelial cells, macrophages and some immune cells (Sarjeant & Stephens, 2012). Expansion of adipose tissue is led by increases in both the number and size of adipocytes (Ali *et al*, 2013). On the one hand, increasing synthesis and storage of triacylglycerols (TAGs) in adipocytes increases the size of these cells, which is defined as hypertrophic adipose tissue growth; on the other hand, differentiation from fibroblast like preadipocytes increases the number of adipocytes, which is referred to as hyperplastic adipose tissue growth and also adipogenesis (Engin, 2017). Adipogenesis includes two phases: first, pluripotent mesenchymal stem cells develop into preadipocytes and thereby become committed to the adipogenic lineage; second, these preadipocytes differentiate into lipid-laden adipocytes. The signaling mechanisms underlying preadipocyte determination and the early stages of adipocyte differentiation remain poorly characterized; thus deciphering these mechanisms could have important implications for the treatment of obesity and related conditions by identifying signaling proteins that could be targeted in people with these disorders. Changes in intracellular calcium (Ca^2+^) constitute a universal signaling mechanism that mediates a variety of important physiological processes (Berridge *et al*, 2003). However, the role of Ca^2+^ signals as mediators of adipocyte differentiation still remains far from clear. Thus pharmacologically induced Ca^2+^ changes have been shown to both stimulate and inhibit adipocyte differentiation *in vitro*, depending on the point in the differentiation process at which such changes were induced (Ntambi & Takova, 1996); however, one potential problem with such approaches is that they are unlikely to mimic the complex spatiotemporal Ca^2+^ signaling patterns that may be occurring during endogenous differentiation. In terms of downstream effectors of Ca^2+^ signals during adipocyte differentiation, the Ca^2+^-activated protease calpain has been implicated in adipogenic differentiation of preadipocyte cells. Thus calpain mRNA levels were shown to be increased during the first few days of adipogenesis, and inhibition of this protease repressed differentiation by preventing c/EBPβ binding to the promoter of the c/EBPα gene (Patel & Lane, 1999). Other studies have suggested that the Ca^2+^ activated kinase CaMKII (Zhang *et al*, 2018), and the Ca^2+^ activated phosphatase, calcineurin (Liu & Clipstone, 2007; Neal & Clipstone, 2002), may both play roles in adipocyte differentiation. However, the source of the upstream Ca^2+^ signal(s) that is regulating these downstream effectors during adipocyte differentiation remains to be identified. One important way in which Ca^2+^ signals are generated is through release of Ca^2+^ from intracellular stores by the action of Ca^2+^ mobilizing second messengers. The three primary Ca^2+^ mobilizing messengers are inositol trisphosphate (IP_3_), cyclic ADP ribose (cADPR), and nicotinic acid adenine dinucleotide phosphate (NAADP). It has been known for some time that IP_3_ and cADPR release Ca^2+^ from the endoplasmic reticulum (ER) via IP_3_ receptors (IP_3_Rs) and ryanodine receptors (RyRs), respectively (Galione & Chuang, 2020; Galione & Churchill, 2000). In contrast, NAADP targets a Ca^2+^ store located in endolysosome-like acidic organelles (Galione, 2011). Importantly, there appears to be a complex interrelationship between these three signaling pathways, with NAADP acting to induce local Ca^2+^ release from acidic organelles that can then act as a ‘trigger’ to stimulate global Ca^2+^ release from the ER via IP_3_ Rs and RyRs (Galione & Churchill, 2002), but also recent evidence indicating that this relationship is bidirectional. Thus, we have recently shown that Ca^2+^ released from the ER can activate the NAADP pathway both by stimulating Ca^2+^-dependent NAADP synthesis and by activating NAADP-regulated Ca^2+^ channels (Morgan *et al*, 2013). The identity of such NAADP-regulated Ca^2+^ channels remains controversial. In line with their assumed location in acidic organelles, studies by ourselves and others have identified the two pore channels (TPCs; gene names *Tpcn1* and *Tpcn2*) as endolysosomal ion channels and integral components of the intracellular receptor for NAADP (Brailoiu *et al*, 2009; Calcraft *et al*, 2009a; Ruas *et al*, 2010; Jin *et al*, 2020; Ruas *et al*, 2015). Moreover, gene knockout (KO) of TPC1 and TPC2 in mouse embryonic fibroblasts abolished their ability to mediate NAADP-regulated, Ca^2+^ signaling responses (Ruas *et al*, 2015). In this study, we investigated the role of TPC2 during adipogenesis, both by studying adiposity in TPC2 KO mice, and by investigating the pattern of expression and functional role of TPC2 during adipocyte differentiation *in vitro* and by investigating the association between genetic variants at *TPCN2* and body fat distribution in humans.

## Results

### TPC2 KO mice of both sexes have abnormal adiposity compared to wild type controls

To determine whether there were any differences in fat or lean content of *Tpcn2* KO mice, mice of both sexes of this strain were analysed using time domain nuclear magnetic resonance (TD-NMR), at 3, 6 or 9 months of age. Male *Tpcn2* KO mice were significantly less heavy than wild type (WT) controls at all ages studied, mainly due to lower fat mass. Lean mass was not significantly different in *Tpcn2* KO males, except in the 9 month old cohort, in which significantly reduced lean mass was observed for *Tpcn2* KO males. This translated into an overall reduced body fat percentage with concomitant increase in percentage of lean composition (Fig. S1A). *Tpcn2* KO females were also significantly lighter than WT controls, up to 6 months, and similar to the male group this was due to reduced fat mass. The reduced fat mass and normal lean mass seen in 3-and 6-month-old females resulted in a decreased body fat percentage and increased percentage in lean composition. In contrast, the 9-month *Tpcn2* KO female group showed no significant differences in any of the parameters measured, when compared to the WT controls (Fig. S1B). To further investigate the physiological condition of *Tpcn2* KO mice, an intraperitoneal glucose tolerance test (IPGTT) was carried out (Fig. S2). No differences in the response to a glucose challenge were detected between *Tpcn2* KO mice and WT controls.

### TPC2 mRNA expression is up-regulated transiently during adipocyte differentiation in vitro

Differences in adiposity in TPC2 KO mice *in vivo* might have a number of different underlying causes. In this study we decided to focus on whether TPC2 might be involved in one specific process linked to adiposity, namely adipocyte differentiation. To see whether TPC2 might play a role in adipocyte differentiation *in vitro*, we used the C3H10T1/2 cell line, a mesenchymal stem cell line that can differentiate along adipogenic, myogenic and chondrogenic pathways (Taylor & Jones, 1979). In the presence of a cocktail of insulin/insulin-like growth factor 1, dexamethasone and IBMX in serum-containing medium, C3H10T1/2 cells differentiate into adipocytes. Progression of adipogenesis was monitored by Oil Red O staining or AdipoRed fluorescence (Fig. 1A), which both measure accumulation of lipid droplets, and RT-qPCR analysis of mRNA expression of adipogenic markers (Fig. 1B), to ensure that the morphological changes observed in cells subjected to the standard differentiation protocol were indicative of true adipogenesis. Early transient upregulation of *c/Ebpδ* mRNA expression, followed by sustained *Pparγ2* and *Srebp1* mRNA expression and later *aP2* mRNA expression, which is the expected sequential pattern of expression of these genes during adipocyte differentiation, confirmed adipogenesis at the molecular as well as the morphological level (Fig. 1B). We next studied whether *Tpcn2* mRNA expression levels changed during the differentiation process; notably, these increased significantly early during adipocyte differentiation, and in a transient manner, with *Tpcn2* mRNA levels peaking at around 4 hours post-induction and returning to starting levels by around 48 hours after the start of induction (Fig. 1C).

**Figure 1:** TPC2 mRNA expression is up-regulated transiently during adipocyte differentiation in vitro. A AdipoRed and Oil Red O staining of differentiating C3H10T1/2 cells, immediately before start of differentiation induction with adipogenic medium, and 4 and 8 days after the start of treatment. Accumulation of the stain indicates lipid droplet formation. Scale bar: 100μm B mRNA levels of adipogenic marker genes in differentiating C3H10T1/2 cells. (a) *Pparγ2*; (b) *Srebp1*; (c) *aP2*; (d) *c/Ebpδ*. n=2 (*c/Ebpδ*); n=3 (*Pparγ2, aP2, Srebp1*). C *Tpcn2* mRNA levels in differentiating C3H10T1/2 cells. n=3. D C3H10T1/2 differentiation medium components. (FCS: fetal calf serum) Data information: In (B), data are presented as mean ± SEM, values are normalised to the geometric mean of 18S and ubiquitin c mRNA expression and presented as fold-change relative to 0h value. §p<0.05; §§p<0.01 (ANOVA, post-hoc Dunnett test vs 0h). In (C), data are presented as mean ± SEM, §§p<0.01 (ANOVA, post-hoc Dunnett test vs 0h)

### Tpcn2 mRNA expression during adipocyte differentiation is up-regulated by a cAMP signal

To investigate which components of the adipocyte differentiation cocktail were responsible for the observed changes in *Tpcn2* mRNA expression, C3H10T1/2 cells were treated with the individual components separately, and harvested at 4 hours post-induction, a stage at which significant change in expression was previously seen. *Tpcn2* mRNA levels were significantly increased by IBMX, to a level comparable to that achieved by complete differentiation medium, whilst the other components of the cocktail had no significant effect on *Tpcn2* expression (Fig. 2A). Moreover, treatment with medium lacking IBMX but containing all other components of the differentiation cocktail, failed to induce *Tpcn2* mRNA expression (Fig. 2B). This finding suggested a potential involvement of cAMP signaling in the induction of *Tpcn2* expression, since IBMX inhibits cAMP phosphodiesterase (PDE), and thus acts to increase cAMP levels (Thompson, 1991). To further confirm the involvement of the cAMP signaling pathway in the induction of *Tpcn2* gene expression, alternative methods of raising levels of cAMP were used to investigate whether this is the signaling intermediate by which IMBX affects *Tpcn2*. Rolipram, like IBMX, is a cAMP PDE inhibitor; however, while IBMX is relatively nonspecific and may affect cGMP as well as cAMP levels, rolipram is a specific inhibitor of PDE IV, thus it specifically raises cAMP levels (Thompson, 1991). In contrast, forskolin increases cAMP levels by activating the synthesising enzyme adenylyl cyclase (Laurenza *et al*, 1989). We found that both rolipram and forskolin increased *Tpcn2* mRNA levels after 4 hours treatment (Fig. 2C). The increase for both drugs was similar to, though slightly lower than, that seen with IBMX. Further evidence for a role of cAMP in mediating IBMX-induced *Tpcn2* mRNA upregulation was provided by our finding that pretreatment with the adenylyl cyclase inhibitor SQ22536 significantly reduced this IBMX-induced rise in expression (Fig. 2D) (Haslam *et al*, 1978).

**Figure 2-.** *Tpcn2* mRNA expression during adipocyte differentiation. A Determination of the component(s) of the adipogenic differentiation medium that stimulates *Tpcn2* mRNA expression. n=3. Grey bar: sample at 4 hours, n=1. B Effect of IBMX on *Tpcn2* mRNA expression in 2-day post-confluent C3H10T1/2 cells. n=3. C Expression of *Tpcn2* mRNA in C3H10T1/2 cells in response to drugs that alter cAMP levels. Cells were treated for 4 hours at 2 days post-confluence with GM (growth media) containing 50μM forskolin (in DMSO), 100μM rolipram (in DMSO), 500μM IBMX (in 100% ethanol) or vehicle. n=3. D Effect of the adenylyl cyclase inhibitor SQ22536 on IBMX-induced induction of *Tpcn2* mRNA expression. 2-day post-confluent cells were preincubated for 30 minutes with GM containing 300μM SQ22536 or vehicle (water) before 500μM IBMX was added. n=3. E, F, G Comparison of *Tpcn2* mRNA in non-treated (control) and (E) PKI (PKA inhibitor) / (F) ESI-09 (Epac inhibitor) / (G) 666-15 (CREB inhibitor) treated C3H10T1/2 cells. At 2 days post confluence, 10 μM PKI / 25 μM ESI-09/ 1 μM 666-15 were added to the medium during the entire time course of differentiation. n=5-8, Data information: Data are presented as mean ± SEM. In (A-D), values are normalised to the geometric mean of *18S* and *ubiquitin c* mRNA expression and presented as fold-change relative to GM condition. In (A), ###p<0.001, (ANOVA, post-hoc Tukey test). In (B), ##p<0.01; ###p<0.001, (ANOVA, post-hoc Tukey test). In (C), **p<0.01; ***p<0.001 (student’s t-test). In (D), ##p<0.01; ###p<0.001, (ANOVA, post-hoc Tukey test). In (E-G), values were normalized to the geometric mean of *β-Actin* mRNA expression and are presented as expression relative to 0hr. **p<0.01 (student’s t-test).

Our finding that the cAMP signaling pathway is involved in the regulation of *Tpcn2* gene expression during adipocyte differentiation raises the question of which proteins that are downstream of cAMP in this pathway mediate these effects. Previous studies have shown that cAMP can activate protein kinase A (PKA), or alternatively exchange protein activated by cAMP (Epac) (Cheng *et al*, 2008). To investigate the possible involvement of PKA and Epac in the induction of *Tpcn2* gene expression, we studied whether inhibitors of these proteins had an impact on such expression. We found that PKI, an inhibitor of PKA, significantly inhibited *Tpcn2* mRNA expression 2 hours after induction of adipocyte differentiation (Fig. 2E). Treatment with ESI-09, an Epac inhibitor (Ahmed *et al*, 2019), significantly inhibited *Tpcn2* mRNA expression at 2 hours and this effect continued to the 4 hour time point (Fig. 2F). PKA and Epac can both mediate gene expression via the transcription factor cAMP-response element binding protein (CREB) (Kelly, 2018). To study whether CREB plays a role in the cAMP-induced expression of the *Tpcn2* gene, we investigated the effects of a CREB inhibitor, 666-15 (Xie *et al*, 2019). We found that this significantly reduced the expression of *Tpcn2* mRNA during adipocyte differentiation (Fig. 2G). Our findings suggest that both PKA and Epac may be involved in the cAMP-mediated regulation of *Tpcn2* gene expression, via a mechanism involving CREB.

### TPC2 plays a functional role during adipocyte differentiation in vitro

The striking transient upregulation of *Tpcn2* mRNA expression during differentiation of C3H10T1/2 cells into adipocytes *in vitro*, suggests that TPC2 may be playing a functional role in this process. We investigated this possibility initially by ectopically expressing recombinant TPC2 protein in C3H10T1/2 cells; thus, we used a lentiviral vector to express mCherry-tagged TPC2, with mCherry alone used as a control. Expression of mCherry or mCherry-tagged TPC2 in infected cells was confirmed by immunoblotting with an anti-mCherry antibody (Fig. 3A), and by analysis of mCherry fluorescence and anti-mCherry immunofluorescence. (Fig. 3B; Fig. S3) The effects of heterologous TPC2 expression on adipogenic differentiation was assessed by measuring AdipoRed fluorescence (Fig. 3C). Lentiviral-mediated expression of TPC2 did not affect adipogenic differentiation of C3H10T1/2 cells in response to the standard differentiation protocol. However, when IBMX was omitted from the differentiation medium, this led to almost complete suppression of adipogenesis, but if TPC2 was expressed virally, cells exhibited significantly higher levels of differentiation than mCherry-infected controls (Fig. 3C). In addition, preadipocyte cells were isolated from WT or *Tpcn2* KO mice and induced to differentiate into adipocytes. While cells isolated from WT mice showed a high degree of differentiation, those from *Tpcn2* KO mice were inhibited in their differentiation capacity to a significant degree (Fig. 3D).

**Figure 3-.** TPC2 plays a functional role during adipocyte differentiation *in vitro*. A Anti-mCherry western blot confirming expression of lentivirally-delivered mCherry-tagged TPC2 in C3H10T1/2 cells (a); anti-mCherry western blot confirming expression of lentivirally-introduced mCherry (b). B Confocal images of mCherry fluorescence (red) and anti-mCherry immunofluorescence (green) in C3H10T1/2 cells expressing lentivirally-introduced mCherry and mCherry tagged TPCs. Scale bar: 30μm. C AdipoRed quantification of adipogenesis in C3H10T1/2 cells lentivirally infected with mCherry or mCherry-tagged TPC2. Cells were treated according to standard differentiation medium protocol (complete DM); with standard protocol omitting IBMX (DM-IBMX); or maintained in growth medium (GM), and were assayed after 8 days. n=8-12. D AdipoRed quantification of adipogenesis in primary preadipocyte cells from WT and *Tpcn2* KO mice treated with standard differentiation protocol and control group maintained in growth medium, assayed after 8 days. n=6. Data information: In (C, D), data are presented as mean ± SEM. **p<0.01, ***p<0.001 (student’s t-test).

### TPC2 regulation of adipocyte differentiation is mediated by calpain

Our finding that TPC2 appears to be playing a functional role in adipocyte differentiation *in vitro* raises the question of the mechanism by which it does this. Given the evidence implicating TPC2 as an NAADP-regulated Ca^2+^ channel, one possibility is that a Ca^2+^-regulated effector protein downstream of TPC2 is involved in the differentiation of preadipocytes into adipocytes. Calpain is a Ca^2+^-dependent cytosolic protease that binds to membranes on its activation and proteolyses substrate proteins in a specific manner; a previous study showed that calpain is required for differentiation of 3T3-L1 preadipocytes into adipocytes (Patel & Lane, 1999). Interestingly, this same study showed that calpain expression is induced by changes in cAMP levels, just as we have shown in this current study for TPC2. To investigate whether calpain activity is regulated by Ca^2+^ signals mediated by TPC2, we studied changes in calpain activity during differentiation of primary preadipocytes obtained from WT or *Tpcn2* KO mice, into adipocytes. For the WT preadipocytes, what appears to be a gradual rise in calpain activity after induction of differentiation up to 8 hours, is followed by an approximately three-fold spike in activity at 24 hours, after which activity decreases and levels off by 4 days (Fig. 4A). In contrast, for the *Tpcn2* KO preadipocytes, we saw a gradual increase in calpain activity that continues up to 8 days, but no sign of a transient burst of activity at around 24 hours as was seen for the WT controls (Fig. 4B). This indicates that calpain activity during adipocyte differentiation is substantially reduced by *Tpcn2* KO, in a qualitatively distinct manner, and suggests that calpain is an effector of TPC2 in this process.

**Figure 4-.** Calpain activity during adipocyte differentiation. A, B Variation in calpain activity in the process of primary preadipocyte differentiation. (A) WT preadipocytes. n=8. (B) *Tpcn2* KO preadipocytes. n=3-8. Data information: In (A, B), data are presented as mean ± SEM. ###p<0.001 (ANOVA, post-hoc Tukey test, compared between consecutive time points).

### CREB phosphorylation during adipocyte differentiation is regulated by TPC2

Our finding that *Tpcn2* mRNA expression is regulated by cAMP/PKA/Epac/CREB implies that TPC2 is downstream of CREB during adipocyte differentiation. However, there are often reciprocal interactions between components in signaling pathways. We therefore decided to investigate whether *Tpcn2* KO had any effect on the active, phosphorylated state of CREB during adipocyte differentiation, using an ELISA assay to compare levels of total, and phosphorylated, CREB, during differentiation of primary preadipocytes from WT or *Tpcn2* KO mice, into adipocytes. This analysis showed that compared to a steadily increasing phosphorylation level of CREB in WT preadipocytes, CREB phosphorylation was inhibited in *Tpcn2* KO preadipocytes after approximately 1-2 days following induction of differentiation, although the *Tpcn2* KO preadipocytes had an initially higher level of phosphorylated CREB than WT preadipocytes up to day 1 of the differentiation process (Fig. 5B). Ca^2+^/calmodulin-dependent protein kinase (CaMK) triggers phosphorylation of CREB by complex mechanisms (Aimes *et al*, 2000; Ma *et al*, 2014). To investigate whether TPC2 affects the phosphorylation of CREB via a mechanism involving CaMK, we studied the effects of KN 93, a selective CaMKII inhibitor (Bonnefond *et al*, 2015). We found that treatment with this inhibitor leads to an inhibition of CREB phosphorylation, particularly after day 2 of differentiation, in a way that partially mimicks the effects of *Tpcn2* KO (Fig. 5C,D). This suggests that TPC2 may normally be playing a role in enhancing CREB phosphorylation during adipocyte differentiation, via a mechanism involving CaMKII (Fig. 7).

**Figure 5-.** CREB phosphorylation during adipocyte differentiation. A, B, C Levels of CREB and phosphorylated CREB during preadipocyte differentiation. (A) WT preadipocytes. n=3. (B) *Tpcn2* KO preadipocytes. n=3. (C) WT preadipocytes treated with KN 93. D Ratio of phosphorylated CREB to total CREB during adipocyte differentiation. Data information: In (A-C), data are presented as mean ± SEM. §§§p<0.001 (one-way ANOVA was used with post-hoc Dunnett test comparing all timepoints to expression at 0h).

**Figure 6-.** Genetic variation at the TPCN2 locus is associated with altered fat distribution. A *TPCN2* is most highly expressed in adipose compared with other metabolic tissues. RNA-Seq data from GTex. B *TPCN2* expression in whole and fractionated adipose tissue. RNA-Seq data from 11 females. AT, adipose tissue, mADS, mature adipocytes, SVF, stomovascular fraction, TPM, transcripts per million. ***p ≤ 0.0001, two-tailed paired Student’s t-test. C Locus zoom showing a common intronic SNV at *TPCN2*, rs2305498, is associated with increased waist-hip ratio adjusted for BMI.

**Figure 7-.** Key components modulators in cAMP signaling and adipogenesis. Schematic showing key components with their pharmacological modulators in cAMP signaling pathway and how they regulate adipogenesis. Figure created with BioRender.com. (AC=adenylyl cyclase; PDE= phosphodiesterase; CREB= cAMP-response element binding protein).

### Genetic variation at the TPCN2 locus is associated with altered fat distribution

To determine the relevance of our findings to humans we examined the tissue expression pattern of *TPCN2* in GTEx (GTEx Portal). We found that among metabolic organs and tissues, abundance of *TPCN2* mRNA was highest in adipose tissue (Fig. 6A). Furthermore, based on a survey of RNA sequencing data from fractionated adipose tissue from 11 females, we found that *TPCN2* expression was highest in stromovascular cells compared with mature adipocytes (Fig. 6B). We next interrogated data from the GIANT consortium and UK Biobank combined genome-wide association study (GWAS) meta-analysis (Pulit *et al*, 2019). This analysis revealed that an independent signal at the *TPCN2* locus was associated with waist-hip ratio adjusted for body mass index (WHRadjBMI) specifically in females (beta = 0.0208, p=6.06E-11) (Fig. 6C). No association was detected in males (beta = 0.0019, p=0.6). The lead single nucleotide variation (SNV) at this signal (rs2305498) lies within an intron of *TPCN2* and is in high mutual linkage disequilibrium (LD) with a rs66958567 (r^2^=0.79) which is a strong expression quantitative trait locus (eQTL) for *TPCN2* in visceral (NES = 0.34, p=5.6E-13) and subcutaneous thigh adipose tissue (NES = 0.32, p=8.8E-11) based on data from GTEx (https://gtexportal.org). The same LD block also contains a missense variant in *TPCN2* (rs35264875) although this is not predicted to be deleterious by PolyPhen and SIFT. Notably, the WHRadjBMI increasing allele at this signal is associated with lower adipose *TPCN2* expression consistent with the pro-adipogenic effect of mouse *Tpcn2 in vitro*. Collectively these data suggest that *TPCN2* plays an important and sexually dimorphic role in adipose tissue biology and fat distribution in humans.

## Discussion

In this study we explored for the first time the link between the two-pore channel, TPC2, and adipogenesis. Our interest was initially stimulated by the observation that *Tpcn2* KO mice of both sexes have both a reduced absolute fat mass and a reduced percentage of fat at 3 and 6 months compared to wild-type controls. These findings show that elimination of *Tpcn2* expression has an effect on adiposity in mice. These findings raise the question of how loss of *Tpcn2* expression might give rise to such an effect. One problem in pursuing this issue further mechanistically is that with such a whole animal KO, loss of *Tpcn2* expression may have various effects due to changes in the properties of a number of different cell types/tissues, some of which may oppose each other. In addition, a common phenomenon in mice KOs is the occurrence of compensation for loss of expression of the knocked-out gene by changes in the levels and patterns of expression of other genes. This complicates interpretation of what a KO phenotype may reveal about the normal physiological role of a gene *in vivo*. We thus chose in the remainder of this study to focus on one potential function for TPC2 in the regulation of the related phenomena of adiposity and adipogenesis, by investigating whether TPC2 plays a role in adipocyte differentiation *in vitro*.

This study provides the first demonstration of a specific cell signaling pathway that can affect the expression of a member of the TPC gene family (*Tpcn2* in mice; *TPCN2* in humans). We found that changes in cAMP levels lead to changes in *Tpcn2* mRNA expression and previous studies have shown that cAMP can activate protein kinase A (PKA), or alternatively exchange protein activated by cAMP (Epac), both of which can work via CREB (Cheng *et al*, 2008). Our findings using specific inhibitors of these proteins suggest that both PKA and Epac may be involved in the cAMP-mediated regulation of *Tpcn2* gene expression, and demonstrate a direct role for CREB in this process (Fig. 7).

That TPC2 may play a functional role during adipocyte differentiation, is suggested by two main findings in the current study. First, we show that over-expression of TPC2 in C3H10T1/2 cells can partially overcome the block to adipocyte differentiation due to the absence of IBMX from the differentiation cocktail. It remains to be shown why this ‘rescue’ is only partially effective; one possible explanation could be that IBMX induces other changes that are necessary for differentiation to occur so that TPC2 expression on its own is only partially able to substitute for absence of cAMP signaling. Second, we show that loss of endogenous TPC2 function in primary preadipocytes from *Tpcn2* KO mice substantially inhibits the ability of these cells to differentiate into adipocytes.

Since previous studies by ourselves and others have previously identified TPCs as integral components of the NAADP-regulated Ca^2+^ signaling pathway, we chose to study whether the downstream mediator of TPC2 action during adipocyte differentiation might be calpain, since this Ca^2+^-sensitive protease has previously been shown to be required for differentiation of 3T3-L1 preadipocytes into adipocytes (Patel & Lane, 1999). The fact that *Tpcn2* KO only appears to affect part of the change in calpain activity during adipocyte differentiation, namely that which occurs transiently at around 24 hours after the start of the differentiation process, is interesting given that *Tpcn2* mRNA expression also occurs transiently, but in this case peaks at 4-8 hours after the induction of differentiation. One possible explanation for these findings would be that there are multiple calpain isoforms that are active during the differentiation of preadipocytes to adipocytes, but only one is regulated by Ca^2+^ signals that are generated via TPC2. A previous study suggested that µ-calpain rather than m-calpain was responsible for the control of adipogenesis of ST-13 preadipocytes established from adult primitive mesenchymal cells, through the regulation of the levels of PPARs and C/EBP (Yajima *et al*, 2006). It would be interesting therefore in future studies to investigate whether a specific calpain isoform is being activated by a Ca^2+^ signal originating from TPC2 and whether this isoform is µ-calpain, since this has been specifically implicated in adipocyte differentiation. In addition, we also showed that the CREB phosphorylation level was enhanced by *Tpcn2* KO in the first 24 hours of adipocyte differentiation, although it peaked on day 2 and quickly fell to its base level while in WT preadipocytes, CREB phosphorylation gradually increased during adipocyte differentiation. This suggests that while our findings indicate that TPC2 gene expression is regulated by a mechanism involving cAMP/PKA/EPAC/CREB, there is a reciprocal regulation of CREB activity by TPC2. This raises the question of how TPC2 might be regulating CREB. One possibility, given that CREB can be phosphorylated by the Ca^2+^/calmodulin-dependent kinases CaMKII and CaMKIV (Sun et al., 1994), is that Ca^2+^ signals induced through TPC2 influence CREB phosphorylation via one or both of these kinases. In line with this, we showed in the current study that pharmacological inhibition of CaMKII partially mimicks the effect of *Tpcn2* KO on CREB phosphorylation. Since our findings also indicate that CREB activates *Tpcn2* expression, this suggests a process of positive feedback involving TPC2 and CREB (Fig. 7).

In keeping with the results of the *in vitro* studies, we also demonstrate that the minor allele of a common intronic SNV at *TPCN2*, rs2305498, is associated with increased WHRadjBMI; a surrogate marker of body fat distribution and an independent predictor of cardiometabolic disease risk. Interestingly, the same allele is also associated with reduced *TPCN2* expression in subcutaneous adipose tissue and increased estimated heel bone mineral density. Based on these findings we speculate that *TPCN2* may limit adipose tissue expansion, thereby promoting upper-body fat accumulation, by altering mesenchymal stem cell fate and promoting adipogenesis at the expense of osteoblastogenesis. This hypothesis is consistent with the expression pattern of *TPCN2* in fractionated human adipose tissue and the early and transient induction of *Tpcn2* gene expression during adipogenesis of C3H10T1/2 mesenchymal stem cells. Future, studies should focus on confirming that *TPCN2* is the effector gene and adipose tissue the target tissue of the WHRadjBMI associations at the *TPCN2* locus, as well, as identifying the effector cell type(s) mediating these associations. The role of *TPCN2* on osteoblastogenesis should also be investigated.

## Materials and Methods

### Animals

*Tpcn2* KO (homozygote *Tpcn2*^YHD437^) mice, previously established by the Parrington/Galione labs (Department of Pharmacology, University of Oxford) (Calcraft *et al*, 2009b) were used. Strain-matched (C57BL/6) WT mice were used as controls. Mice were killed by cervical dislocation or asphyxiation. All experiments and procedures were approved by the local animal welfare and ethics committee of the University of Oxford, in accordance with UK guidelines for the use of experimental animals.

### TD-NMR scannin

Before scanning, mice carcasses were thawed for 24 hours at 4°C followed by 4 hours at room temperature. These samples were scanned in the same manner as live animals (Halldorsdottir *et al*, 2009). Previous investigations have set a precedent for MR analysis of frozen-thawed mice (Tinsley *et al*, 2004). Mice were weighed then scanned using a Bruker Minispec LF50 Whole Body Composition Analyser. Fat mass and lean mass data were acquired using Bruker Minispec Plus software.

### Culture, treatment and adipogenic differentiation of C3H10T1/2 cells

C3H10T1/2 cells (clone 9, passage 18, cells obtained from American Type Culture Collection (ATCC)) were incubated at 37°C with 5% CO_2_ in Dulbecco’s modified Eagle’s medium (DMEM) (Gibco, 61965) supplemented with 10% fetal bovine serum (FBS) and 1% penicillin/streptomycin, refreshed every second day, and passaged before exceeding 80% confluence. Differentiation was induced at 1-2 days post-confluence, designated ‘day 0’. This involved the growth medium being aspirated and replaced with complete DM (growth medium with addition of 1μM insulin, 0.5mM 3-isobutyl-1-methylxanthine (IBMX) and 1μm dexamethasone). After 2 days, this was replaced with DM/ins, and after another 2 days, this was replaced with base DM. Differentiation was allowed to proceed for a total of 8-10 days, with the base DM refreshed every 2 days. For the untreated control condition, cells were maintained in GM, and refreshed every 2 days.

### Isolation, treatment culture and differentiation of primary preadipocyte cells

Abdominal adipose tissue from mice was obtained at the age of 10-18 weeks. A Preadipocyte Isolation Kit (ab196988, Abcam) was used for preadipocyte isolation and purification according to the manufacturer’s instructions. The preadipocytes were cultured in DMEM/F12, supplemented with 10% fetal bovine serum (FBS) and 1% penicillin/streptomycin, for 48 hours until adherence. The differentiation was induced following the same standard differentiation protocol as used for C3H10T1/2 cells, but replacing DMEM with DMEM/F12.

### Oil red O staining

A solution of 0.2% Oil Red O (w/v) in isopropanol was filtered. Cells were fixed in 4% paraformaldehyde for 30 min, incubated for 5 mins in 60% isopropanol, then incubated in the Oil Red O solution for 10 mins before being imaged using a Leica DM 500B microscope.

### AdipoRed staining and quantification

Cells were processed with AdipoRed (Lonza) reagent according to the manufacturer’s instructions before being observed under a fluorescence microscope. Fluorescence was measured using a Typhoon 9400 Variable Mode Imager, with excitation of 488nm and use of a BP 580/30 emission filter. The photomultiplier tube (PMT) was adjusted to maximize the signal while avoiding saturation, and resolution was 100μm. Fluorescence was quantified using Fiji/Image J software. Data were normalized to an average background of empty wells or, in the case of lentivirally-infected cells, to average reading for cells infected with the relevant lentivirus but not treated with AdipoRed, to control for overlap between mCherry and AdipoRed fluorescence emission.

### Quantitative PCR analysis

Total RNA from C3H10T1/2 cells was isolated and purified using a QIAGEN RNeasy mini kit according to the manufacturer’s instructions. The RNA concentration was checked using a NanoDrop spectrophotometer and integrity was checked by electrophoresis on a 1.5% agarose gel. cDNA was synthesized from RNA samples using a high capacity cDNA reverse transcription kit (Applied Biosystems) according to the manufacturer’s instructions. The cDNA synthesized was stored at −80°C until analyzed. A Roche LightCycler 480 System was used for qPCR. All primers were designed using the Roche Universal Probe Library Assay Design Centre, which also identified suitable fluorescent probes for each primer pair. A reaction mixture of cDNA (0.00075ng/μl-3ng/μl), gene-specific forward and reverse primers (0.1μM each) and Universal ProbeLibrary Probe (0.1μM) in LightCycler 480 Probes Master was prepared. All samples were run in duplicate within each experiment and results averaged. Cycling conditions were 50°C for 2 mins; 95°C for 10 mins; a 30-50x cycle: 95°C for 15 secs; 60°C for 1 min; then an end cycle: 40°C for 30 secs. LightCycler 480 SW 1.5 software acquired data and produced graphs quantifying the fluorescence of each sample plotted against cycle number and calculated the Cp for each sample. The value for each gene of interest was normalized to the averaged housekeeping gene value and were expressed as values relative to a control (0 hour timepoint for differentiation time-course experiments, GM or vehicle control for pharmacological interventions).

### Lentiviral expression of TPC2

Lenti-X HEK293T cells (Clontech) were grown at 37°, 5% CO_2_ in DMEM supplemented with 10% tetracycline-free foetal calf serum (BioSera), 1mM sodium pyruvate, 100U/ml penicillin-G-sodium, 100μg/ml streptomycin sulphate and 2mM L-glutamine. When 50-80% confluent, cells were co-transfected with viral vector and Lenti-X HT packaging mix (Clontech, 15μl per 3μg vector) using a jetPEI system (Polyplus transfection) according to the manufacturer’s instructions. Medium was harvested 48 hours after transfection, filtered through a sterile 0.45μm polyethersulfonate filter (Whatman puradisc 25 AS), and checked for the presence of infectious virons by applying a sample to Lenti-X GoStix (Clontech). When cells reached 50% confluence, mCherry, mCherry-tagged TPC lentiviral medium or control medium was added (1ml/T75 flask) along with polybrene to 4μg/ml final concentration.

### Fluorescence and immunofluorescence imaging

C3H10T1/2 cells (seeded on coverslips) were washed in PBS and fixed for 15 mins in 4% paraformaldehyde, then mounted with ProLong Gold antifade mounting medium (Invitrogen) (for mCherry fluorescence observation) or processed for immunofluorescence: cells were permeabilised in PBS/0.5% Triton X, then blocked in 2% goat serum followed by incubation in 5F8 monoclonal against mCherry, supplied by A. Rottach, University of Munich, diluted 1:500, for 1 hour at room temperature, and incubation in goat anti-rat Alexa Fluor 488 (Invitrogen), 5μg/ml. Coverslips were mounted and slides viewed using a Zeiss Meta 510 confocal LSM.

### Immunoblot analysis

Cells were lysed by RIPA (Sigma) and dissolved in 5 × Laemmli sample buffer (ThermoFisher) supplemented with phosphatase inhibitor cocktails (Sigma). The proteins were boiled at 96 °C for 5 mins, separated by SDS-polyacrylamide gel electrophoresis (SDS-PAGE), and transferred to polyvinylidene difluoride membranes. To block nonspecific binding sites, membranes were incubated in Tris-buffered saline containing 0.1% (v/v) Tween 20 and 5% (w/v) skim milk powder before primary antibodies of the specificity of interest were added. Films were observed by LI-COR Odyssey 9120 Digital Imaging System.

### Calpain activity analysis

Calpain activity of differentiating primary preadipocytes was measured using a Calpain Activity Assay Kit (ab65308) according to the manufacturer’s instructions. Fluorescence was read by a Tecan M200 Pro microplate reader.

### CREB phosphorylation analysis

Cells were subject to same lysis procedure as for Western Blot analysis. CREB levels (total/phosphorylated) were measured using a CREB InstantOne ELISA kit (Invitrogen) according to the manufacturer’s instructions.

### Analysis of TPCN2 genetic variants in humans

Abdominal and gluteal subcutaneous adipose tissue biopsies were obtained from 11 healthy females recruited from the Oxford BioBank (OBB), aged 30-50 years with BMI ranging from 22-27kg/m^2^. The study was approved by Oxfordshire Clinical Research Ethics Committee (08/H0606/107+5) and all subjects gave written informed consent. Stromal vascular fraction (SVF) cells and mature adipocytes were separated from whole adipose tissue biopsies following collagenase (Roche) digestion (1 mg/ml in Hanks’ buffered salt solution) and in the case of SVF cells additional treatment with red cell lysis buffer. Total RNA was extracted as described by Collins et al (Collins *et al*, 2010). RNA integrity was assessed using a 2100 Bioanalyzer (Agilent Technologies) and samples with a RNA integrity (RIN) number of 8 and above were selected for RNASeq analysis. These samples were sent to the the Wellcome Trust Centre for Human Genetics where a quality control report was generated and a library of read data compiled. Differentially expressed genes were identified by comparing the RNAseq profiles between SVF, mature adipocytes and to the whole adipose tissue samples.

### Statistical analysis

All data are presented as +/-standard error of the mean (SEM). For most data comparisons, unpaired 2-tail student’s t-test was used (*p<0.05; **p<0.01; ***p<0.001). For multiple comparison of 3 or more data sets, one-way analysis of variance (ANOVA) was used with post-hoc Tukey test (##p<0.01; ###p<0.001). For statistical analysis of variation in gene expression and calpain activity during adipogenic differentiation time-course, one-way ANOVA was used with a post-hoc Dunnett test comparing all timepoints to expression at time zero (§p<0.05; §§p<0.01). Statistical analyses were conducted using GraphPad Prism8.

## Author Contributions

JP, YZ, LHC and RT designed and performed experiments, analyzed all data, and wrote the manuscript. MR, DG, MT and CC performed experiments and helped to write the manuscript.

## Expanded view

### Tpcn2 KO mice of both sexes have abnormal adiposity compared to wild type controls

**Figure EV1.**
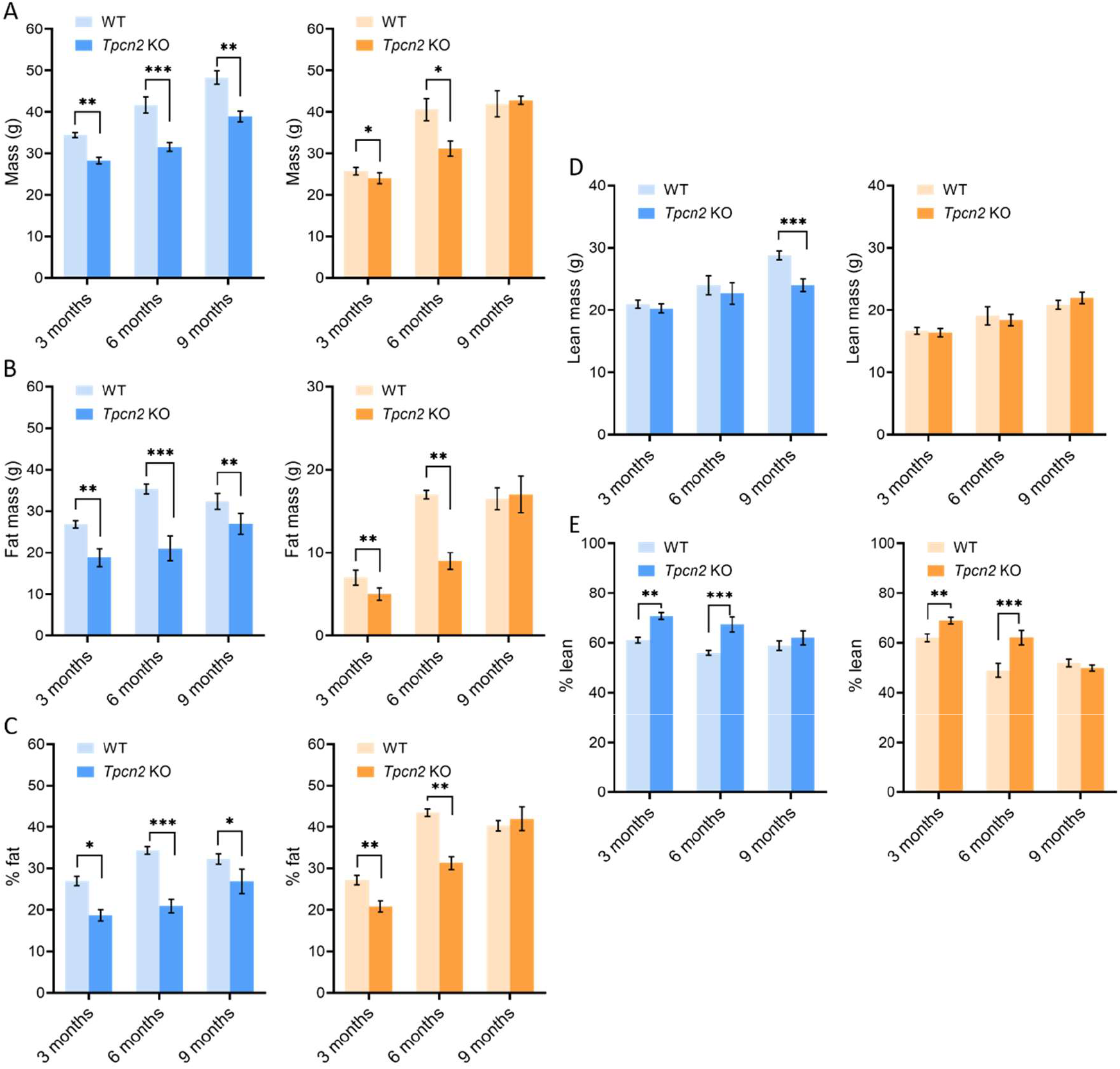
Male (blue) and female (orange) *Tpcn2* KO and WT body composition quantified by TD-NMR: A) total body mass; B) fat mass; C) percentage fat; D) lean mass; E) percentage lean. n=7-8. *p<0.05; **p<0.01; ***p<0.001 (student’s t-test).

### Intraperitoneal glucose tolerance test (IPGTT) of TPC KO mice

In response to a glucose challenge, *Tpcn2* KO mice showed a slight, non-significant tendency towards blood glucose peaking higher than in WTs, with no significant difference in total glycaemia.

**Figure EV2:**
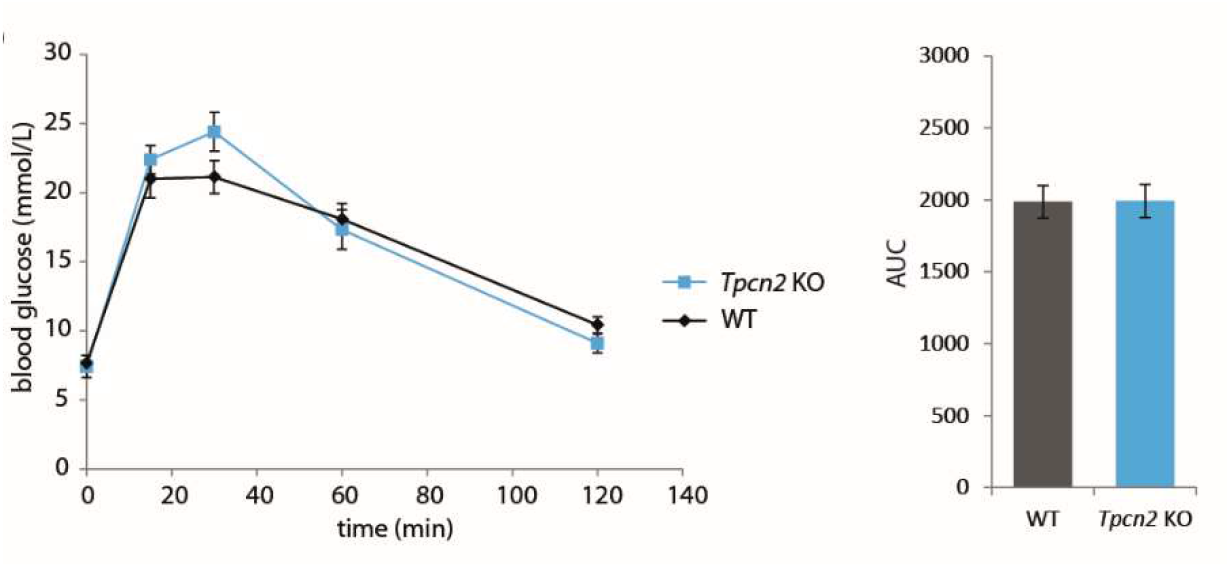
Intraperitoneal glucose tolerance test (IPGTT). Response to 2g/kg IP glucose following overnight fast in male *Tpcn2* KO and strain-matched WT mice aged 9-11 weeks. *Tpcn2* KO (n=11), WT (n=8); student’s t-test. AUC = area under the curve.

### Anti-mCherry immunofluorescence

Figure S2 shows anti-mCherry immunofluorescence in MEFs infected with lentiviruses bearing cDNA encoding mCherry and mCherry-tagged TPC2. Anti-mCherry immunofluorescence was clearly visible in infected cells, corresponding to obvious mCherry fluorescence in mCherry-tagged TPC2 infected cells. The expression of mCherry-tagged TPC2 would be predicted to also be punctate based on the previously observed lysosomal localization of TPC2, but this was less obvious – possibly due to excessive overexpression, or simply to overexposure of the confocal images. mCherry fluorescence in cells infected with mCherry only was less bright and more difficult to distinguish from background but was confirmed as genuine mCherry fluorescence by colocalization with anti-mCherry immunofluorescence.

**Figure EV3:**
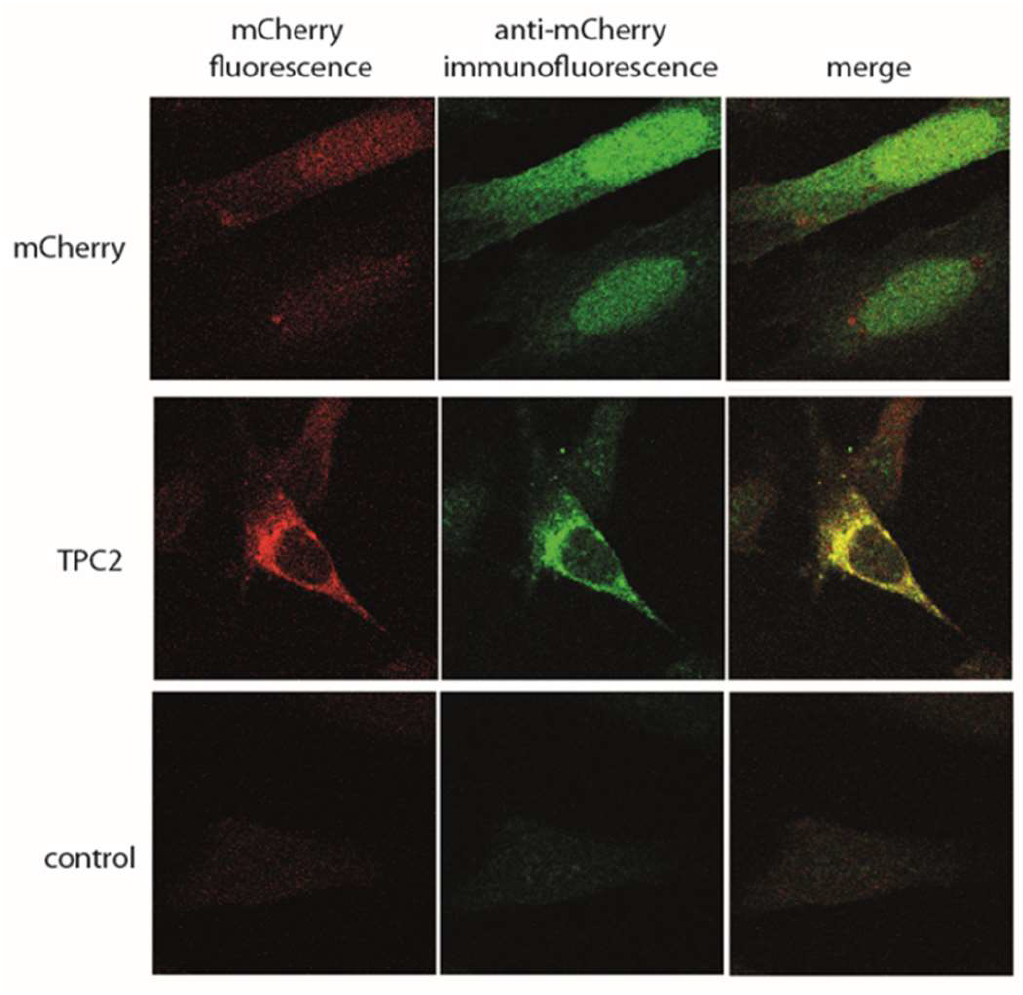
Expression of lentivirally-transduced mCherry and mCherry-tagged TPC2 protein in paraformaldehyde-fixed MEFs. mCherry fluorescence (red); anti-mCherry immunofluorescence (green).

**Table EV1:**
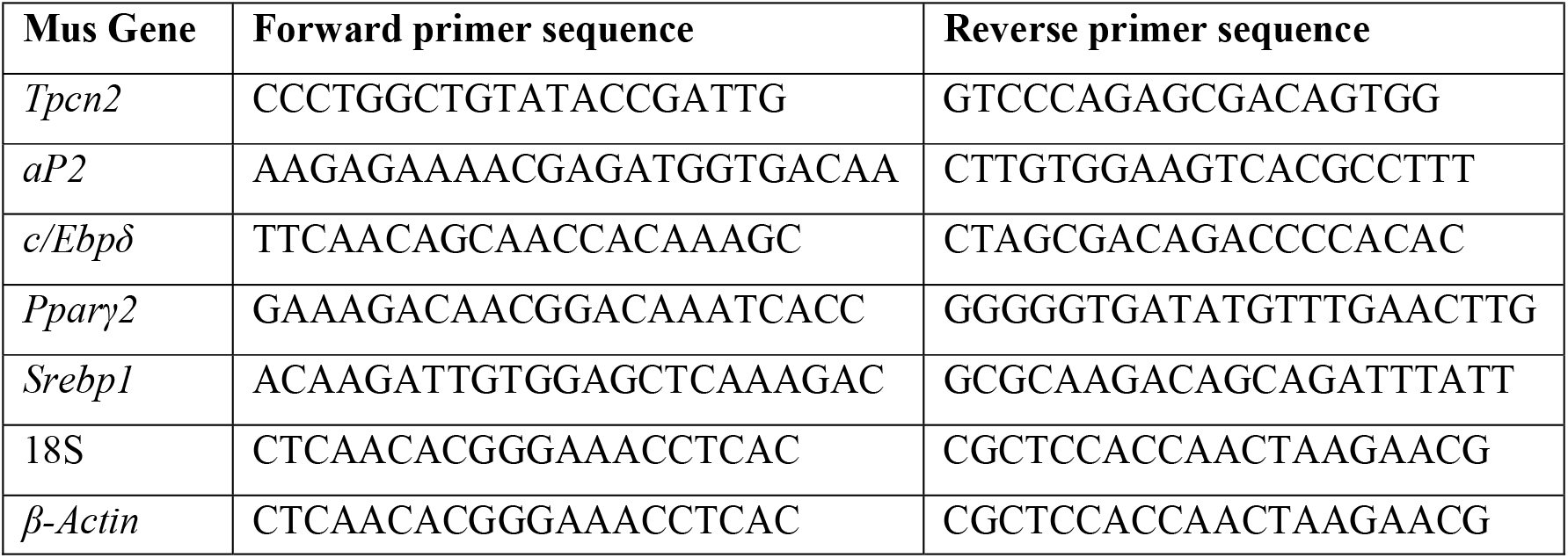
List of oligonucleotides used for qPCR.

